# Comparative risk-assessment of highly pathogenic avian influenza H5 viruses spread in French broiler and layer sectors

**DOI:** 10.1101/2024.09.11.612235

**Authors:** Claire Hautefeuille, Facundo Muñoz, Gwenaëlle Dauphin, Mathilde Paul, Marisa Peyre, Flavie Goutard

## Abstract

Since 2015, French poultry production is threatened almost every year by a reintroduction of highly pathogenic avian influenza H5 viruses. The duck sector was the most concerned by this crisis but other sectors such as broiler, layer and turkey were also affected by outbreaks. The objective of this work was to assess the risk of highly pathogenic avian influenza H5 virus transmission from one farm to another within the French broiler and layer production network.

This study used the WOAH risk assessment framework. After drawing up a scenario tree of virus transmission from one farm to another, data were collected through a literature review or through experts’ elicitation. Three questionnaires were developed according to the experts’ field of expertise: avian influenza, broiler and layer sectors. The experts’ estimates were combined using a beta distribution weighted by their confidence level. A Monte Carlo iteration process was used to combine the different probabilities of the scenario tree and to assess the transmission risk.

In the broiler sector, the highest transmission probabilities were observed if the exposed farm was an indoor broiler farm and the source a broiler farm (indoor or free-range). The high transmission probability between broiler farms integrated within the same association suggests that integration is an important risk factor. Person movement, transport of feed and manure management were the pathways with the highest transmission probabilities between two integrated indoor broiler farms with good biosecurity levels. In the layer sector, the highest transmission probabilities were observed if the source farm was a free-range farm and the exposed farm a production farm (indoor or free-range). The pathways with the highest transmission probabilities were egg transport and person movement. The sensitivity analysis showed that the exposed farm’s biosecurity had a significant impact on the transmission probability.

Our results provide an insight on the role of each type of farms in the virus spread within the French broiler and layer production sectors and will be useful for the implementation of control measures such as movement restriction or vaccination.

## Introduction

Since 2015, France has faced several epizootic waves of highly pathogenic avian influenza (HPAI) viruses [1–3]. The main poultry production sector affected by these waves has been the fattening duck production sector – for the production of duck liver-based delicacies called “foie-gras” [4,5]. During these waves, while the initial introduction of HPAI viruses has likely been due to wild birds [1,6], the large spread of the viruses between poultry farms was linked with the poultry production systems. The structure of the fattening duck sector was one of the main factors facilitating the spread of HPAI [6]. Indeed, in this sector, live birds’ movements occur between the different production stages: rearing, breeding and force-feeding. It has been demonstrated that these movements have played a major role in the spread of HPAI viruses [7,8].

Other poultry production sectors exist in France: the meat sector (broiler, meat duck, turkey, guinea fowl) and layer sector [9]. In the broiler production sector, birds enter the farm as day-old chicks and leave the farm to go to the slaughterhouse. In the layer sector, production is organized in two stages: pullet (from day-old chick to ready-to-lay layers) and layer (from ready-to-lay layers to spent hen). In between these two stages, live bird movements take place to transport ready-to-lay layers from the pullet farm to the layer farm. The same movement pattern is observed for breeders of all species. A live bird movement occurs between the growth stage farm and the fertile adult stage farm. Even if there are no live bird movements between broiler or layer farms, other avian influenza (AI) virus pathways such, as fomite or human movements, may exist [10]. Identified fomite movements may occur through birds picked-up for slaughter, feed delivery, egg collection, manure and litter management, dead bird management and shared equipment [11–15]. Movements involving direct contacts between humans and birds are identified as presenting the higher risk of AI virus spread (veterinarian, staff working on multiple premises, company staff, technical advisors etc.) [16,17].

Few risk assessment studies looked at the impact of the integration level (integrated or independent) or farming conditions (indoor vs. free-range) on the risk of AI viruses introduction in a farm [13,15,18]. Moreover, a few modelling studies collected data on the frequency of fomite and human movements between farms [11,14]. Nevertheless, these studies never took a close look at the origin of these movements, i.e. type of farm from which the movement originated.

So far, no study has been conducted on the risk of AI introduction into a chicken farm from another farm within the chicken production sector in France, even though these sectors have also been strongly affected by outbreaks (329 outbreaks in chicken farms during the 2020-22 period) [2]. The aim of this study was to assess the risk of spread of AI virus within two other French poultry production sectors: broiler and layer. This risk assessment study has been conducted considering different types of farms as described in a previous study [9]. These types of farms were defined according to the sector (broiler and layer)), production type (i.e. future breeders, breeders, production), the farming conditions (i.e. indoor and free-range) and the integration level of the farm. For this purpose, this study assesses the risk of transmission from one farm of a defined type to another farm of the same or different type within the same production sector. It did not consider the transmission risk between farms from different poultry sectors and not the risk of introduction through environmental pathways. The current study focuses on the relative transmission risk, i.e. to identify the type of farms with the highest risk of virus transmission through indirect contacts (human and fomites movements). This study will allow us to identify pathways and farm types most at risk of AI virus transmission.

## Materials and methods

### Risk assessment model

This study focused on the assessment of the risk of introduction described by the WOAH framework [19,20]. The aim of the study was to evaluate the relative risk of viral spread between different types of farms. We assumed that once the virus is introduced into the flock, the probability of a bird being infected, as well as the risk of exposure, is close to 100%, and we did not assess the consequences of AI spread. We considered it highly likely that bird movements from an infected farm would introduce the virus and cause an infection on a free farm. Consequently, we focused solely on the risk of introduction through indirect pathways within a poultry production network. Four pathways were identified in a previous literature scoping review [10]: human with a direct contact with birds; trucks transporting live birds; trucks transporting feeds, manure and dead birds; shared equipment in direct contact with birds (Figure 1). Two other pathways that are known to represent a very low risk were not considered: human and shared equipment with no direct contact with birds or manure [11,15,21]. Environmental pathways such as air-born or rodent as mechanical vector transmission pathways were assumed homogeneous across farm types and excluded from the study. Geographical proximity between farms was not considered either.

**Figure 1:**
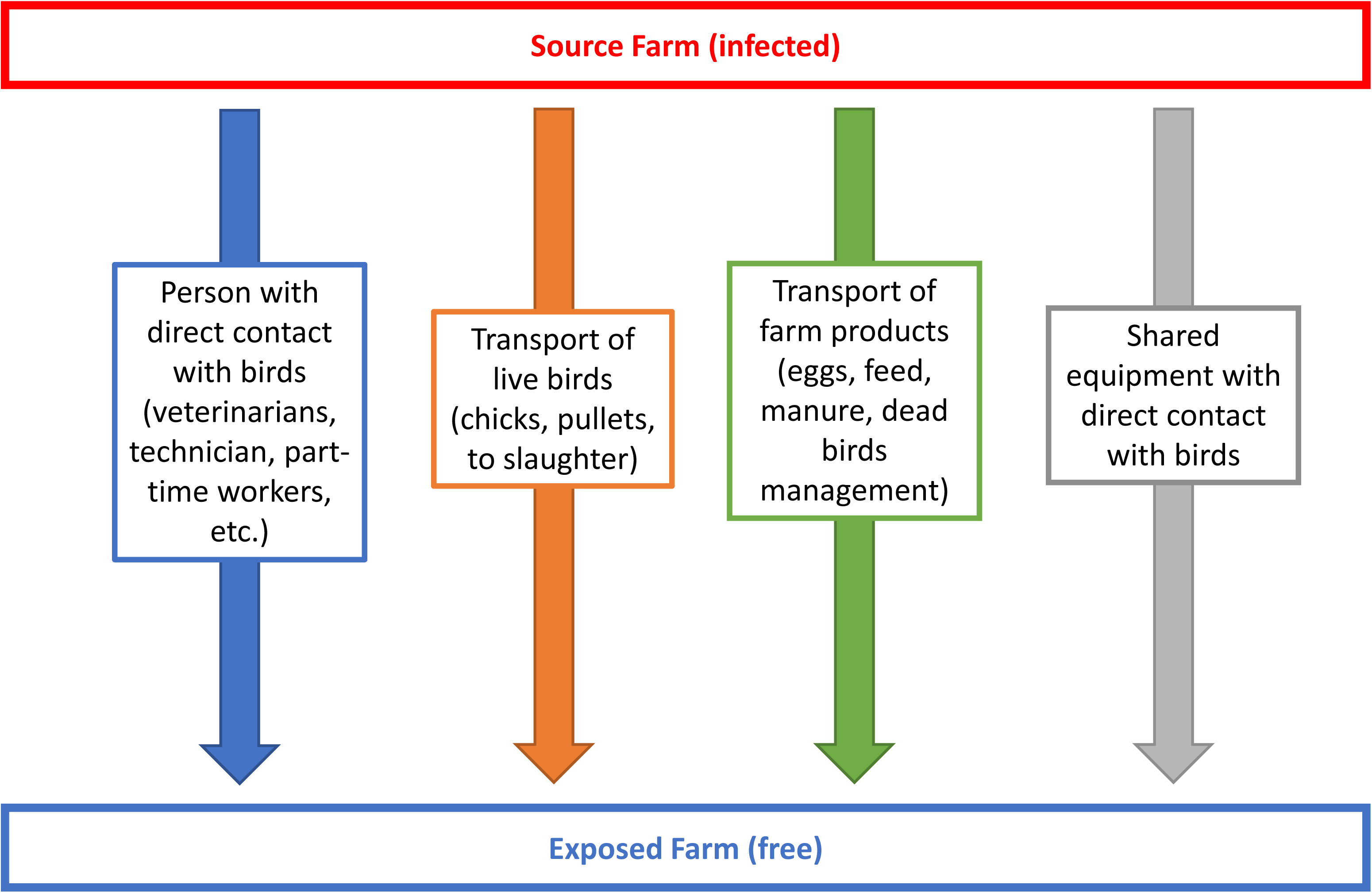
The transmission pathways considered in this risk-assessment study.

The assessment was conducted for the broiler and layer sectors on several types of farms defined in Table 1 and adapted from the description provided by Hautefeuille et al. [9]. Considering the low number of breeder farms and future layers (pullets) farms, we did not include the integration level for these types of farms.

**Table 1:** Listing of the different types of farms studied according to the production type, the farming conditions and the integration level of farms in the French broiler and layer production sector

| Farming type | Production type | Integration level |
| --- | --- | --- |
| Indoor | Broiler | integrated in the same farmers' association |
|  |  | integrated in another farmers' association |
|  |  | independent |
| Free-range | Broiler | integrated in the same farmers' association |
|  |  | integrated in another farmers' association |
|  |  | independent |
| Selection | Breeder | - |
|  | Future breeder | - |

### Data source

The probability of virus introduction from each pathway was estimated using scenario trees as recommended by EFSA [22]. Two scenario trees were built (Figure 2): one to estimate the probability of the non-detection of an infected farm (Figure 2 and Table 2) and one to estimate the probability of viral transmission from the source farm to the exposed farm through a given pathway (Figure 3 and Table 3). The first step of this study was to identify, in the scenario trees, the branches for which estimates of risk probabilities could be obtained from the French legislation. The second step was to generate data for each pathway on the transmission risk, the frequency of occurrence for one type of farm and the probability that this pathway exists between two different types of farms. An expert elicitation survey was conducted to obtain the data not available from the scientific literature.

**Figure 2:**
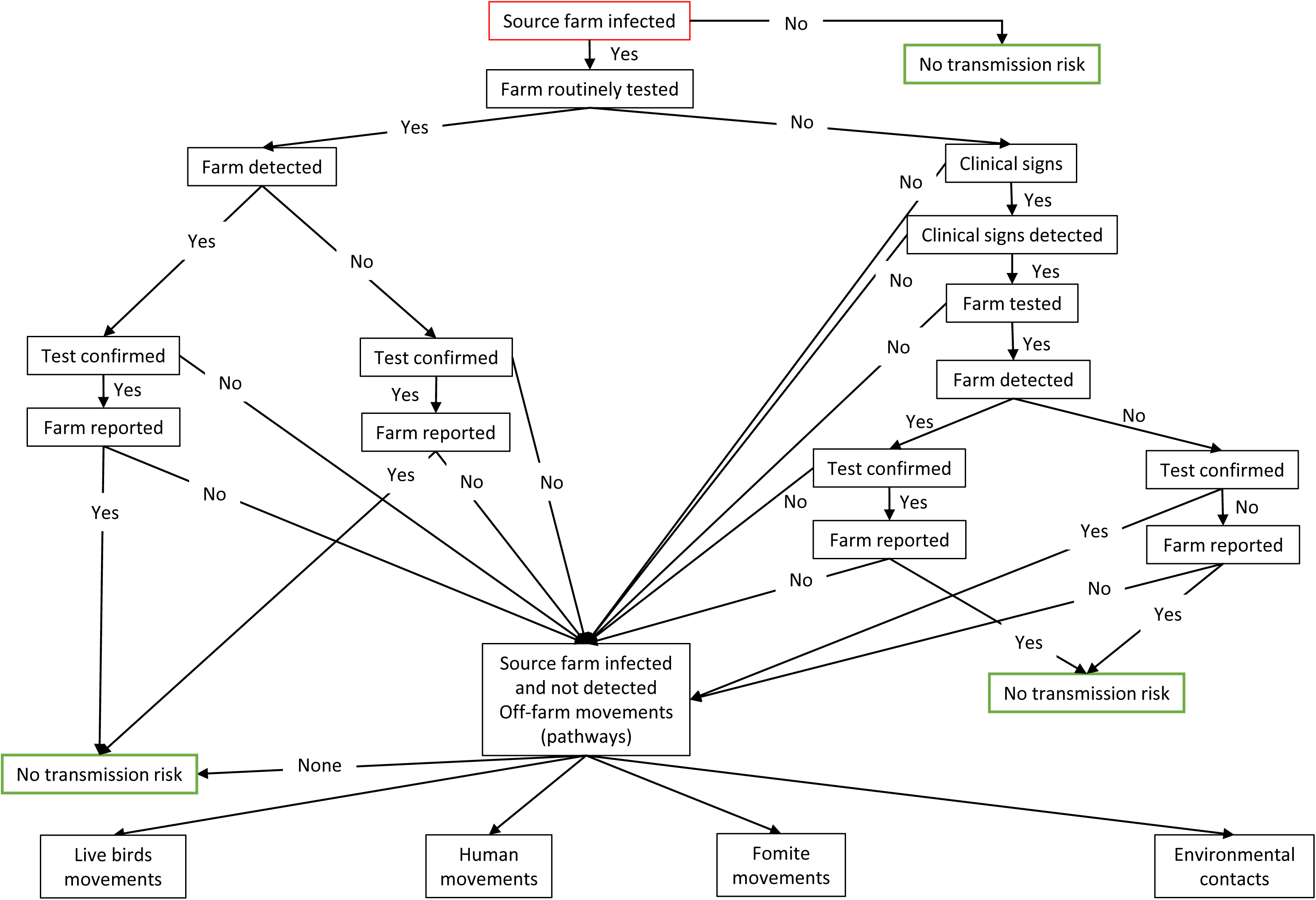
Scenario tree to assess the risk of not detecting an infected farm

**Figure 3:**
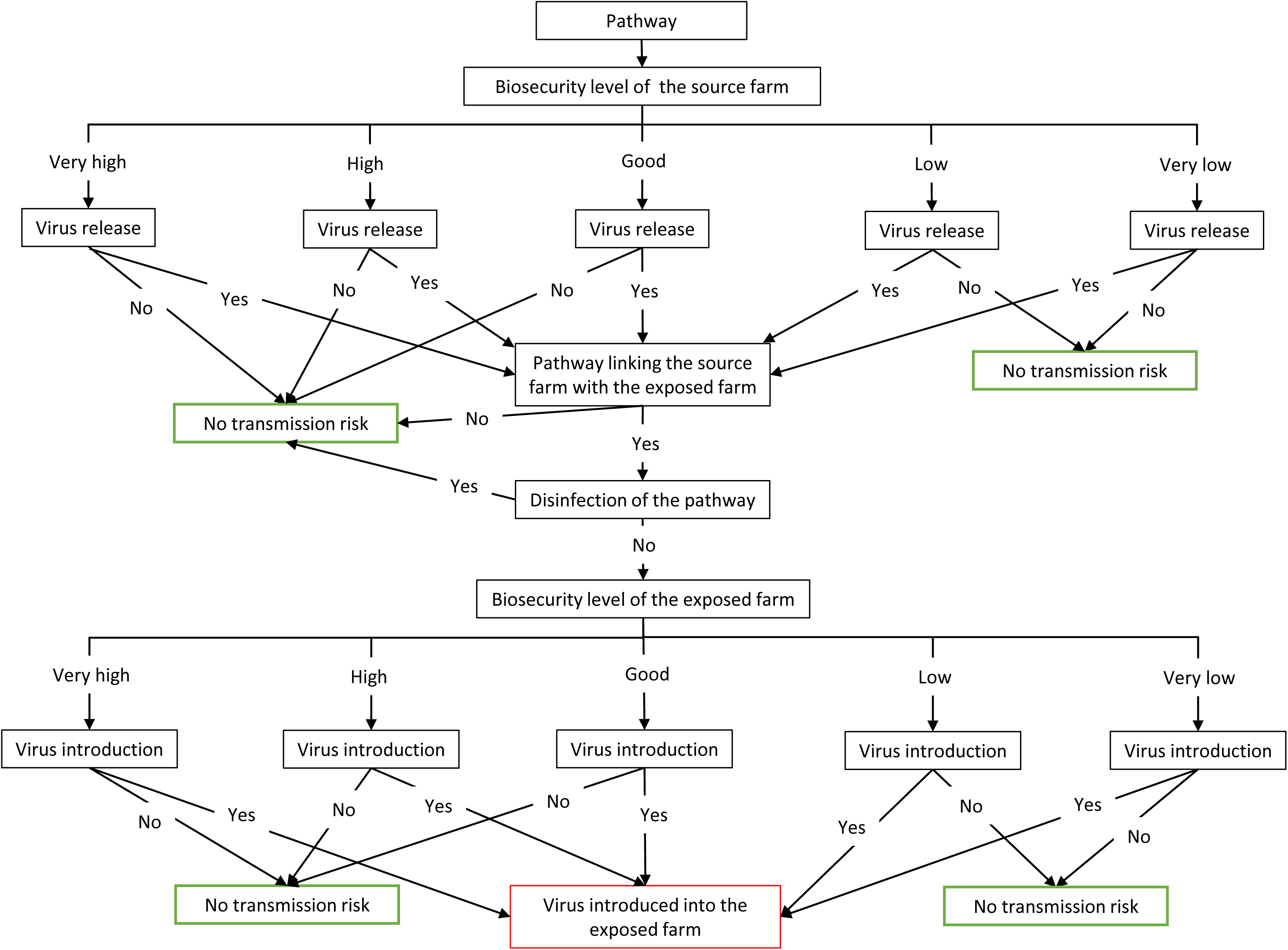
Scenario tree to assess the risk of virus transmission from an infected farm (source) to a free farm (exposed) through a given pathway.

**Table 2:** Probability that an infected farm (source) is not detected

| Node | Parameters estimates | Branch of node | Data sources or justification |
| --- | --- | --- | --- |
| Farm routinely tested | Probability that a farm is routinely tested against HPAI | Yes<br>No | French legislation [23] |
| Time of infection | Estimation of the proportion of farms infected for more or less than 3 days | Less than 3 days<br>More than 3 days | Incubation period [24] |
| Clinical signs | Probability that a trained farmer detects clinical signs of HPAI infection before 3 days of infection | Yes<br>No | Expert elicitations |
|  | Probability that a trained farmer detects clinical signs of HPAI infection after 3 days of infection | Yes<br>No | Expert elicitations |
| Farm tested if clinical signs detected | Probability that a farm is tested if clinical signs are detected | Yes<br>No | Expert elicitations |
| Farm reported if tested positive | Probability that a farm is reported if tested positive | Yes<br>No | French legislation [23] |

**Table 3:** Probability that a silently-infected farm (source farm) transmits the virus to a free farm (exposed farm) for a given pathway

| Node | Parameters estimates | Branch of node | Data sources or justification |
| --- | --- | --- | --- |
| Biosecurity level of source farm (infected farm) | Estimation of the proportion of farms of the same type of farms than source farm in the different biosecurity levels | Very low<br>Low<br>Good<br>High<br>Very High | Expert elicitations |
| Virus release from source farm (infected farm) | Probability that the virus is released from source farm according to the studied pathway | Yes<br>No | Expert elicitations |
| Link between the source farm and the exposed farm | Probability that the studied pathway occurred in the exposed farm | Yes<br>No | Expert elicitations or production cycle values |
|  | Probability that the studied pathway went during the previous week in the source farm | Yes<br>No | Expert elicitations |
| Biosecurity level of exposed farm | Estimation of the proportion of farms of the same type of farms than the exposed farm in the different biosecurity levels | Very low<br>Low<br>Good<br>High | Expert elicitations |
| (free of infection) |  | Very High |  |
| Virus introduction in the exposed farm | Probability that the virus is introduced in the exposed farm according to the studied pathway | Yes<br>No | Expert elicitations |

Fomite (i.e. person, truck or material) disinfection outside farms was considered to be similar for all types of farms and not sufficient to stop the virus spread.

Once the virus is introduced in the exposed farm (free of infection), it was considered that the probability that the virus enters in contact with birds varies according to the pathway. A farm consists of several areas to meet biosecurity criteria as explained in the introduction of the biosecurity grid: public area for visitors, professional area where only the persons and vehicles authorized to work on the farm can go, and the production unit (i.e. the building where the birds are raised with a possible outdoor area). For all studied pathways, the area into which the virus is introduced (Table 4 and Figure 4) and the probability that a virus is transmitted from one area to another (Table 5) were defined.

**Figure 4:**
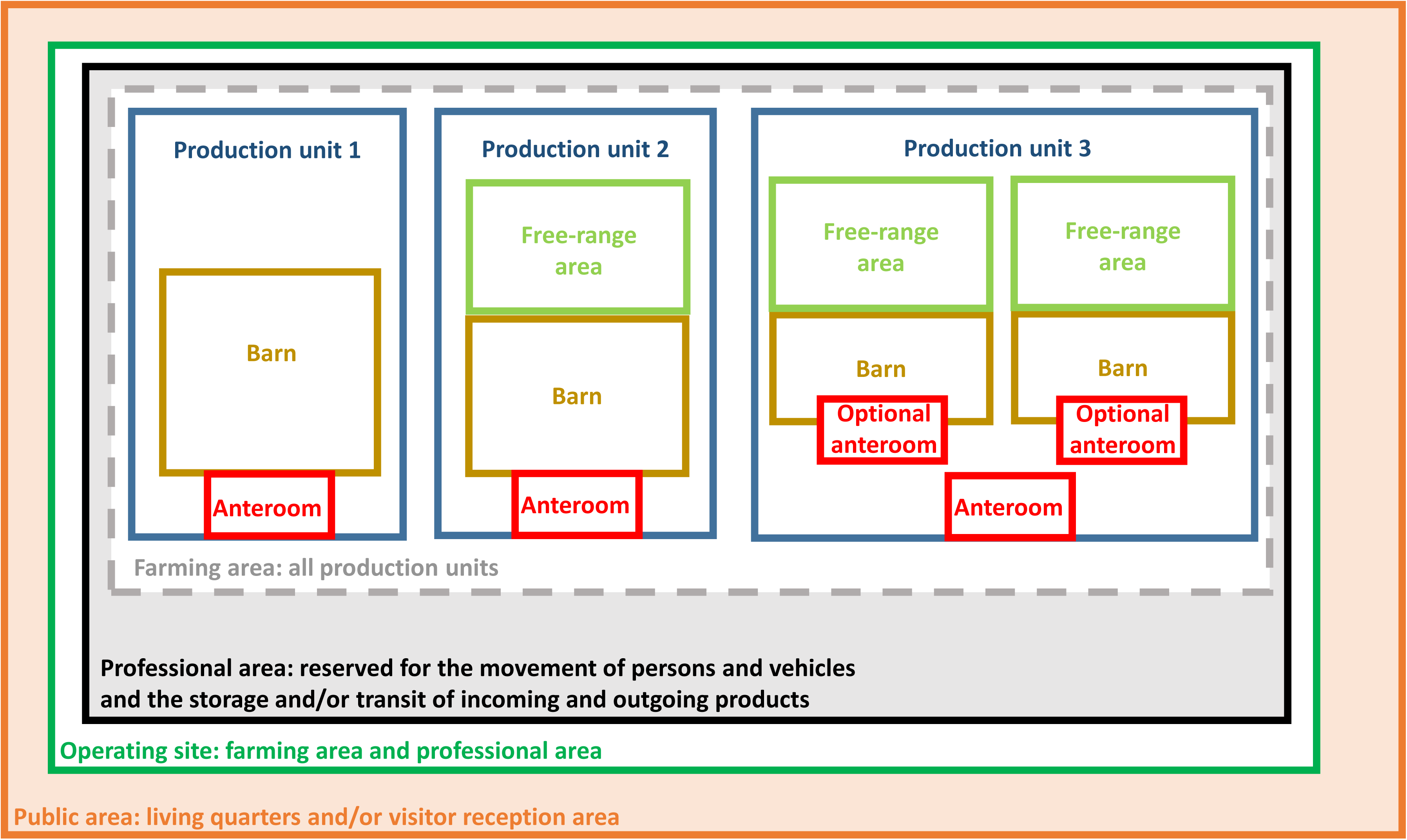
Schematic representation of the different areas composing a broiler farm (based on[25])

**Table 4:** List of the different studied pathways and farm area of virus introduction according to this pathway

| Pathways | Farms concerned | Area of the farm where the virus is introduced |
| --- | --- | --- |
| Human with a direct contact with birds | All | Direct contact with birds |
| Chick transport | Broiler<br>Future breeder |  |
| Birds' pick-up transport (pullet and to slaughter) | All | Professional area |
| Feed delivery | All |  |
| Manure management | All |  |
| Dead bird management | All | Public area |
| Shared equipment | All | Production unit |
| Egg | Layer, Breeder | Professional area |

**Table 5:** Probability that a virus coming into contact with birds after introduction into a free farm of a given biosecurity level

| Node | Parameters estimates | Branch of node | Data sources or justification |
| --- | --- | --- | --- |
| Virus presents in public are | Probability that the virus is introduced in the professional area | Yes<br>No | Expert elicitation |
| Virus presents in professional area | Probability that the virus in introduced in the production unit | Yes<br>No | Expert elicitations |
| Virus presents in production unit | Probability that the virus enters in contact with birds | Yes<br>No | Expert elicitations |

### Experts’ elicitation survey

The aim of this survey was to collect data on the transmission risk for each type of farms (8 farm types for broiler and 7 for layer production) and for each pathway (9 different pathways). We created questionnaires for three different profiles: one for experts on AI epidemiology and poultry farm biosecurity, one for experts on French broiler sector and one for experts on French layer sector.

Experts were recruited according to their experience on the different questionnaire topics. The survey on AI was initially sent to scientists from national French research centres who are currently working and publishing on AI in France. Surveys about the broiler and the layer sectors were initially sent to people who represent these industries at a national level. Following this initial recruitment, a “snowball” recruitment was conducted, in which experts who participated in the survey were asked to recommend other experts to fill out the questionnaire. Experts filled the survey independently without being in contact with each other. The survey was not anonymous so that experts could be contacted for clarification, but all their answers were analysed and presented anonymously.

The questionnaires were designed and deployed online using the platform Survey Monkey (www.surveymonkey.com). For each questionnaire, a pilot survey was sent to two experts in the corresponding field in order to test and optimize the survey. To make the questionnaires shorter, some simplifications were decided and validated by these two experts (Table 6).

**Table 6:** Simplification made according to the probability and the pathway concerned and justification validated by two experts of the poultry production sector.

| Probability concerned | Pathway concerned | Simplification made | Justification |
| --- | --- | --- | --- |
| Probability of virus introduction and release | Pullet transport<br>Birds to slaughter transport | Merge of the two pathways as pick-up transport | Pick-up of birds is conducted in a similar way for pullet and birds to slaughter. |
| Probability that the studied pathway went during the previous week in farm A (i.e. Probability of pathway origin) | Manure<br>Dead birds | Merge of the two pathways | Management similar for both pathways |
|  | Shared equipment | No consideration of the integration level of farms | Management similar regardless the integration level |

The experts were emailed a description of the study objectives and of what was expected for their participation with the link to the online survey. The beginning of the online survey included instructions and contact information of the administering researchers. The questionnaires are available (Supplementary file 1).

Each questionnaire was designed to take about one hour to be filled by experts. Questionnaires started with a description of the experts and questions on their expertise level. The first questionnaire on AI epidemiology was divided in four sections: risk of AI virus introduction in a farm, risk of AI virus release from an infected farm, risk associated with wild birds, and risk of an infected farm going undetected. To simplify the questionnaire, risk was assessed per farm biosecurity level rather than farm type. Grids usually used for biosecurity evaluation process are developed to assess the biosecurity at one farm level in a very detailed way and in the aim to improve the biosecurity of this one farm. Thus, it was necessary to develop a specific less detailed biosecurity grid to order farms within one farming type between five biosecurity levels defined from very low to very high. This biosecurity grid was built based on previous works on the biosecurity evaluation process [7,26] and guidelines developed by the French poultry technical institute (ITAVI) [25] (Supplementary file 2). This grid was validated by two experts outside of the list of experts identified for the risk assessment and currently working on French poultry farms biosecurity. For the five defined biosecurity levels, experts were asked about the probability of introduction or release of the virus for each risk pathway (poultry production related or wild birds related). At the end, the experts were asked to define the probability that an infected farm would go undetected. The other two questionnaires were designed to collect data on broiler and layer sectors. To assess the biosecurity level of a studied farm type, experts were asked to distribute 100 farms of this type between the five different levels in order to have the percentage of farms per biosecurity level (Supplementary file 1). The second section aimed to identify for each introduction pathway and for each type of farm, the frequency that this pathway occurred and the probability that this pathway links the studied farm with another type of farms. The questionnaire concluded on the capacity of a farmer to detect an AI infection based on the type of farm. Regarding the probabilities of virus introduction or release according to the several pathways, experts were asked for the most probable, along with the minimum and maximum values. For question concerning probabilities of pathway occurrence, experts were asked for a range of probabilities (Null: 0%, Negligeable: >0 – 20%, Low: 20-40%, Moderate: 40-60%, High: 60-80% and Very high: >80%). For question concerning frequencies, experts were asked to select on possibility from a range between “more than seven times a week” and “less than once every two months”. For each question, experts were asked to provide their confidence level with a score from 1 to 5 (Table 7) and they could add comments. Finally, for the three questionnaires, the experts were asked some open-ended questions regarding factors that might have influenced their answers. All the answers provided by the experts are available in Supplementary file 1.

**Table 7:** Scoring of confidence levels

| Level of confidence | Score |
| --- | --- |
| Certain | 5 |
| Very confident | 4 |
| Moderately confident | 3 |
| Not confident | 2 |
| Uncertain | 1 |

### Data analysis

Data collected from the SurveyMonkey platform were saved in a database template developed in Microsoft Excel® version 2007. The data from these questionnaires were used to elicit Beta distributions characterising the ensemble of experts’ assessments on each probability of interest. For each probability of interest, the parameters of the Beta distribution were determined in terms of expected value and variance of the distribution, which were in turn calculated from the expert’s responses as follows. For the question asking for the most probable, minimum and maximum values of the target probabilities, the expected value was set at the average of the most probable values weighted by the expert confidence level and the variance was calculated using a formula (1) balancing a maximal variance V_m_ and an average variance V_i_ between experts.

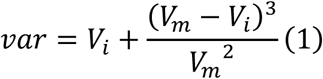

The maximal variance V_m_ corresponds to the variance of a Uniform distribution with support over the maximal interval (i.e. the smallest interval containing all the intervals reported by the experts). The average variance V_i_, by contrast, is the average of the variances of the Uniform distributions over each of the reported intervals. This variance calculation allows the representations of divergent opinions without reducing compatible assessments. For questions requiring only one interval of values (i.e. 21% to 40%, 41% to 60%, etc.), such as questions on between farm contact probabilities, the centre of the interval was considered as the most probable, and the boundaries as the minimum and the maximum value for each expert. The analysis was conducted with R (version 4.0.2).

The frequency of occurrence and the origin of some pathways did not need to be assessed by experts as these values depend on the production cycle. The egg transport was considered to enter into a given breeder farm every day and the possible origin of this truck was always breeder farms. The frequency of chick and pullet delivery and slaughter take off was defined by the production cycle of the different types of production. As it appears that experts could not provide an occurrence frequency for shared equipment, it was arbitrarily considered to occur once a month and validated with the experts who validated the questionnaire.

A Monte Carlo iterative process was carried out on the scenario trees described above, in order to combine the different probabilities and to calculate the transmission probability from one type of farm to another. For each iteration, the probability of realisation of the event for all branches of the scenario tree were randomly sampled from the Beta distribution previously elicited and all probabilities were multiplied to obtain the transmission probability. The simulation process used 50 000 iterations.

In addition, we estimated the relative transmission probabilities between farm types with respect to the pair that exhibited the highest mean transmission probability. The target pair of source and exposed farm types was deemed to be at very high transmission risk when the sampled transmission probability was greater than that of the reference pair in 40% or more of the Monte Carlo iterations. For percentages between 25 and 40%, the pair was considered to be at high transmission risk. For percentages between 15 and 25%, 5 and 15% and below 5%, the pair was considered to be at medium, low and very low transmission risk respectively.

### Sensitivity analysis

A sensitivity analysis was conducted using the one-at-time method applied on the biosecurity level of the source and the exposed farms and on the pathways [27]. This analysis was applied only for two types of farms with the most extreme daily transmission probabilities: one representing the highest and one the lowest. This sensitivity analysis is based on the distribution of the transmission probabilities on 10 000 iterations.

## Results

### Experts’ description

At the end of the process, six experts responded to the questionnaire on AI epidemiology: three had more than ten years of experience, one had over five years and two had more than three years. Seven experts, all with over ten years of experience, responded to the questionnaire on the broiler sector. Five experts, each with over ten years of experience, answered the questionnaire on the layer sector.

### Probability of avian influenza virus transmission

The survey results for the broiler production network indicated that transmission probabilities were highest either from indoor integrated production farms (i.e. not selection farms) to indoor broiler farms (regardless of integration level), or from indoor or free-range production farms to independent indoor farms. These farm pairs were considered to be at very high transmission risk (Table 8). One of the highest median transmission probabilities was observed between independent indoor broiler farms, which served as the reference for relative comparison. The lowest probabilities were found either from selection farms to free-range farms (regardless of the level of integration) or when the exposed farm was a selection farm. High transmission probabilities were also found from free-range or independent indoor farms to integrated indoor farms, and between free-range independent farms. Finally, while transmission probability was of medium level from production to free-range farms, it was low from selection to indoor farms.

**Table 8:** Daily transmission probabilities of HPAI H5 virus between an infected source farm and an exposed farm (free of infection) according to the type of farms in the French broiler production network (median, Q25 and Q75 values of 50 000 iterations). Percentage of probabilities below the one estimated between independent indoor broiler farms: Dark brown: 50-60% (very high risk of transmission), brown: 60-75% (high risk), medium brown: 75-85% (medium risk), light brown: 85-95% (low risk), no colour: over 95% (very low risk).

| Values in % |  | Exposed farm (free of infection) |  |  |  |  |  |
| --- | --- | --- | --- | --- | --- | --- | --- |
|  |  | Indoor broiler |  | Free-range broiler |  | Selection broiler |  |
| Source farm (infected) |  | Integrated | Independent | Integrated | Independent | Breeder | Future breeder |
| Indoor broiler | Integrated in the same farmers' association | 1.2 [0.5 - 3] | 1.0 [0.4 - 2.5] | 0.2 [0.1 - 0.4] | 0.3 [0.1 - 0.6] | 0.004 [0.0005 - 0.02] | 0.001 [0.00008 - 0.01] |
|  | Integrated in another farmers' association | 0.8 [0.3 - 2] | 1.0 [0.4 - 2.5] | 0.2 [0.1 - 0.5] | 0.3 [0.1 - 0.6] | 0.004 [0.0005 - 0.02] | 0.001 [0.00008 - 0.01] |
|  | Independent | 0.6 [0.2 - 1.4] | 1.2 [0.5 - 2.8] | 0.2 [0.1 - 0.5] | 0.3 [0.2 - 0.7] | 0.005 [0.0006 - 0.03] | 0.001 [0.00009 - 0.01] |
| Free-range broiler | Integrated in the same farmers' association | 0.5 [0.2 - 1.5] | 1.0 [0.4 - 2.4] | 0.3 [0.2 - 0.8] | 0.3 [0.1 - 0.7] | 0.004 [0.0005 - 0.03] | 0.001 [0.00008 - 0.01] |
|  | Integrated in another farmers' association | 0.5 [0.2 - 1.5] | 1.0 [0.4 - 2.4] | 0.3 [0.1 - 0.6] | 0.3 [0.2 - 0.7] | 0.004 [0.0006 - 0.03] | 0.001 [0.00009 - 0.01] |
|  | Independent | 0.6 [0.3 - 1.7] | 1.2 [0.5 - 2.8] | 0.3 [0.1 - 0.5] | 0.4 [0.2 - 1] | 0.005 [0.0007 - 0.03] | 0.001 [0.0001 - 0.01] |
| Selection broiler | Breeder | 0.04 [0.009 - 0.2] | 0.06 [0.01 - 0.3] | 0.01 [0.003 - 0.04] | 0.02 [0.005 - 0.05] | 0.005 [0.0005 - 0.03] | 0.001 [0.00008 - 0.01] |
|  | Future Breeder | 0.04 [0.008 - 0.2] | 0.06 [0.01 - 0.3] | 0.01 [0.002 - 0.04] | 0.01 [0.004 - 0.04] | 0.002 [0.0002 - 0.02] | 0.002 [0.0001 - 0.02] |

For the layer production network, the transmission probabilities were the highest between production farms (pullet, indoor or free-range layer) (Table 9). One of the highest median transmission probabilities was between independent free-range layer farms, which were used as the reference farms for the relative risk comparison. Transmission probabilities were the lowest between selection farms. Finally, transmission probabilities were medium from production to pullet farms, from pullet to production farms and from indoor to production farms (except for independent indoor farms).

**Table 9:** Daily transmission probabilities of HPAI H5 virus between an infected source farm and an exposed farm (free of infection) according to the type of farms in the French layer production network (median, Q25 and Q75 values of 50 000 iterations). Percentage of probabilities below the one estimated between independent free-range layer farms: Dark brown: 50-60% (very high risk of transmission), brown: 60-75% (high risk), medium brown: 75-85% (medium risk), light brown: 85-95% (low risk), no colour: over 95% (very low risk)

| Values in % |  | Exposed farm (free of infection) |  |  |  |  |  |  |
| --- | --- | --- | --- | --- | --- | --- | --- | --- |
|  |  | Pullet | Indoor layer |  | Free-range layer |  | Selection layer |  |
|  |  |  | Integrated | Independent | Integrated | Independent | Breeder | Future breeder |
| Source farm (infected) |  |  |  |  |  |  |  |  |
| Pullet |  | 0.2 [0.08 - 0.5] | 0.1 [0.04 - 0.3] | 0.1 [0.05 - 0.3] | 0.1 [0.05 - 0.3] | 0.1 [0.04 - 0.3] | 0.006 [0.001 - 0.03] | 0.003 [0.0005 - 0.02] |
| Indoor layer | Integrated | 0.1 [0.06 - 0.4] | 0.2 [0.09 - 0.6] | 0.2 [0.08 - 0.5] | 0.2 [0.08 - 0.6] | 0.2 [0.09 - 0.6] | 0.006 [0.001 - 0.03] | 0.004 [0.0006 - 0.02] |
|  | Independent | 0.1 [0.06 - 0.4] | 0.2 [0.07 - 0.5] | 0.3 [0.1 - 0.7] | 0.2 [0.09 - 0.6] | 0.2 [0.09 - 0.6] | 0.006 [0.001 - 0.03] | 0.004 [0.0006 - 0.02] |
| Free-range layer | Integrated | 0.2 [0.09 - 0.5] | 0.3 [0.1 - 0.8] | 0.3 [0.1 - 0.9] | 0.4 [0.2 - 0.9] | 0.3 [0.1 - 0.8] | 0.008 [0.002 - 0.03] | 0.005 [0.0008 - 0.02] |
|  | Independent | 0.2 [0.07 - 0.5] | 0.3 [0.1 - 0.7] | 0.3 [0.1 - 0.8] | 0.3 [0.1 - 0.7] | 0.4 [0.1 - 0.9] | 0.008 [0.001 - 0.03] | 0.005 [0.0008 - 0.02] |
| Selection layer | Breeder | 0.008 [0.002 - 0.03] | 0.006 [0.001 - 0.02] | 0.007 [0.002 - 0.03] | 0.007 [0.002 - 0.02] | 0.008 [0.002 - 0.03] | 0.006 [0.0007 - 0.04] | 0.003 [0.0004 - 0.02] |
|  | Future breeder | 0.007 [0.001 - 0.03] | 0.006 [0.001 - 0.02] | 0.006 [0.001 - 0.02] | 0.006 [0.002 - 0.02] | 0.006 [0.002 - 0.02] | 0.003 [0.0004 - 0.02] | 0.004 [0.0005 - 0.02] |

### Sensitivity analysis

The sensitivity analysis was conducted on two farm types: one with the highest transmission probability (integrated indoor broiler farm, Figure 5 and Figure 6) and one with the lowest (future breeder farm for broiler production), both as source (infected) and exposed (free) farms (Supplementary file 3). In both cases, the biosecurity level has a strong influence on transmission probability: it was high when both source and exposed farms had low biosecurity and low when both had high biosecurity (Figure 5).

**Figure 5:**
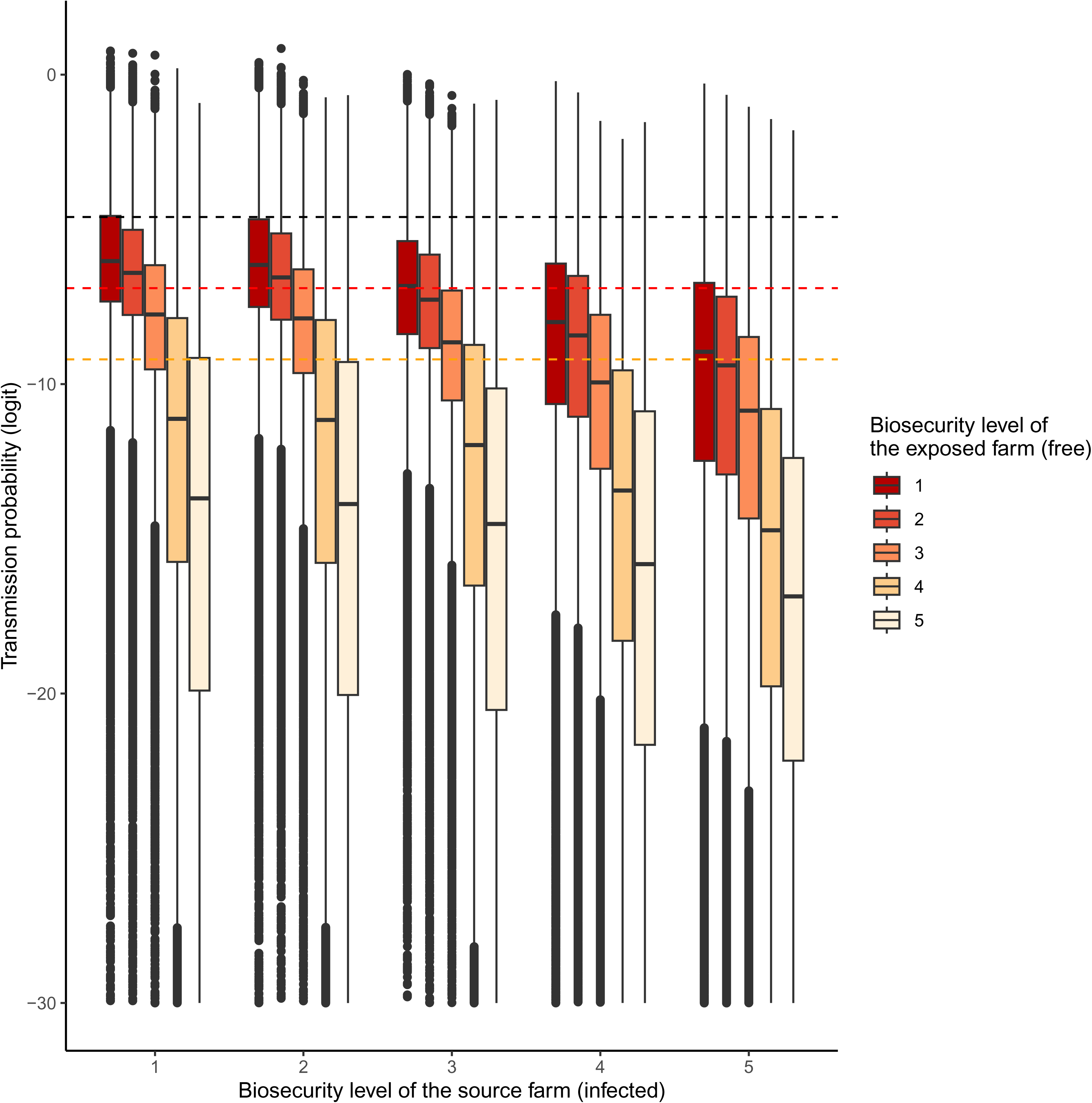
Distribution of the transmission probabilities across all **pathways** according to the biosecurity level of the source farm (x axis) and to the biosecurity level of the exposed farm (colours) with integrated indoor broiler farm as exposed and source farms. Threshold at 1/100 (black), 1/1000 (red) and 1/10 000 (orange).

**Figure 6:**
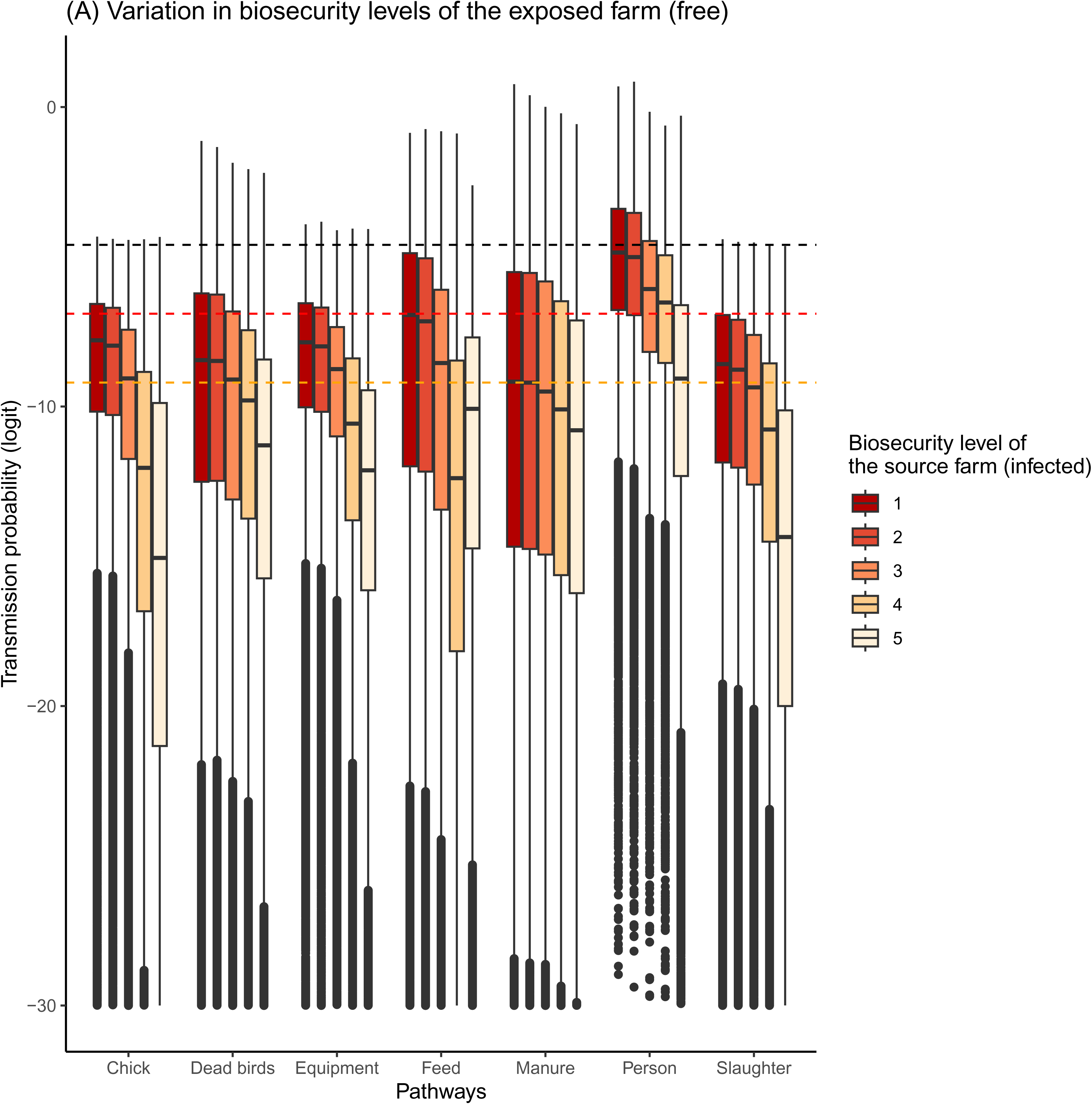
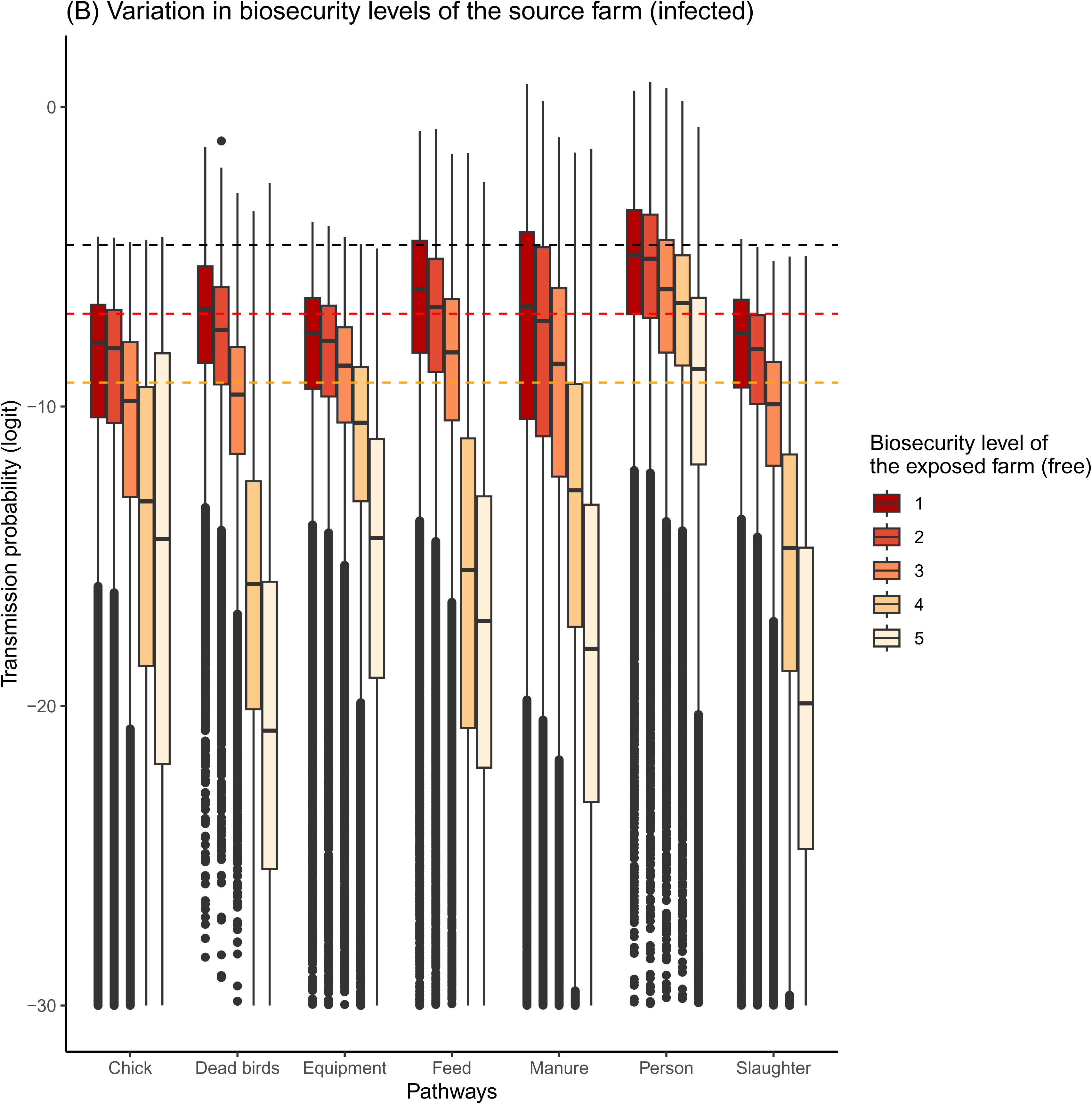
Distribution of the transmission probabilities (A) across all **biosecurity levels of the exposed farm (free)** according to the pathways (x axis) and to the biosecurity level of the source farm (infected) (colours) and (B) across all **biosecurity level of the source farm (infected)** according to the pathways (x axis) and to the biosecurity level of the exposed farm (free) (colours) with integrated indoor broiler farm as exposed and source farms. Threshold at 1/100 (black), 1/1000 (red) and 1/10 000 (orange).

The biosecurity level of the exposed farm had a greater influence than that of the source farm. For all pathways, variations in the source farm’s biosecurity (infected) had low impact on transmission probabilities (Figure 6 A). In contrast, variations in the biosecurity level of the exposed farm significantly influence transmission probabilities (Figure 6 B). Indeed, probabilities were high when the exposed farms had a very low to good level (from 1 to 3 out of 5), and very low when the level was high (above 4), for all pathways except person movement.

### Farm biosecurity and pathways in the broiler sector

Most integrated broiler farms (indoor and free-range) were scored with a good biosecurity level (median value of 45% of indoor and 37% of free range) (Supplementary file 4). Independent indoor farms were scored with a higher biosecurity level (42% at good level) than independent free-range broiler farms (30% at good level). In both systems, independent broiler farms were estimated to have lower biosecurity than integrated farms. Among integrated indoor broiler farms with good biosecurity, the highest transmission probabilities were linked to person movement and transport of feed (median value of 0.23% and 0.03% respectively). These pathways were the most frequent with median occurrence frequency of 21% for person movement and 30% for transport of feed. When considering an integrated indoor broiler farm as an exposed farm, the probability of contact through person and feed movement was high if the source farm belongs to the same farmers’ association, medium if from another farmers’ association and low otherwise (Figure 7). Integrated free-range broiler farms had the lowest transmission probabilities as exposed farm despite similar biosecurity scores than integrated indoor broiler farms. The pathways with the highest transmission probability between two integrated free-range broiler farms at good biosecurity level were material, feed and manure movement (respectively median value of 0.02% and 0.01%). Person movement had less impact on the probability of transmission in free-range farms than in indoor farms, associated with a lower occurrence frequency. For both integrated indoor and free-range farms, transport of birds from hatchery (chicks) or to slaughter and dead bird management had the least impact on transmission probabilities. Movement of birds from hatchery and to slaughter had the lowest frequency of occurrence (associated with the production cycle). Dead bird management was considered to have a high frequency of occurrence, but this pathway is the only one that stops in the public area of the farm.

**Figure 7:**
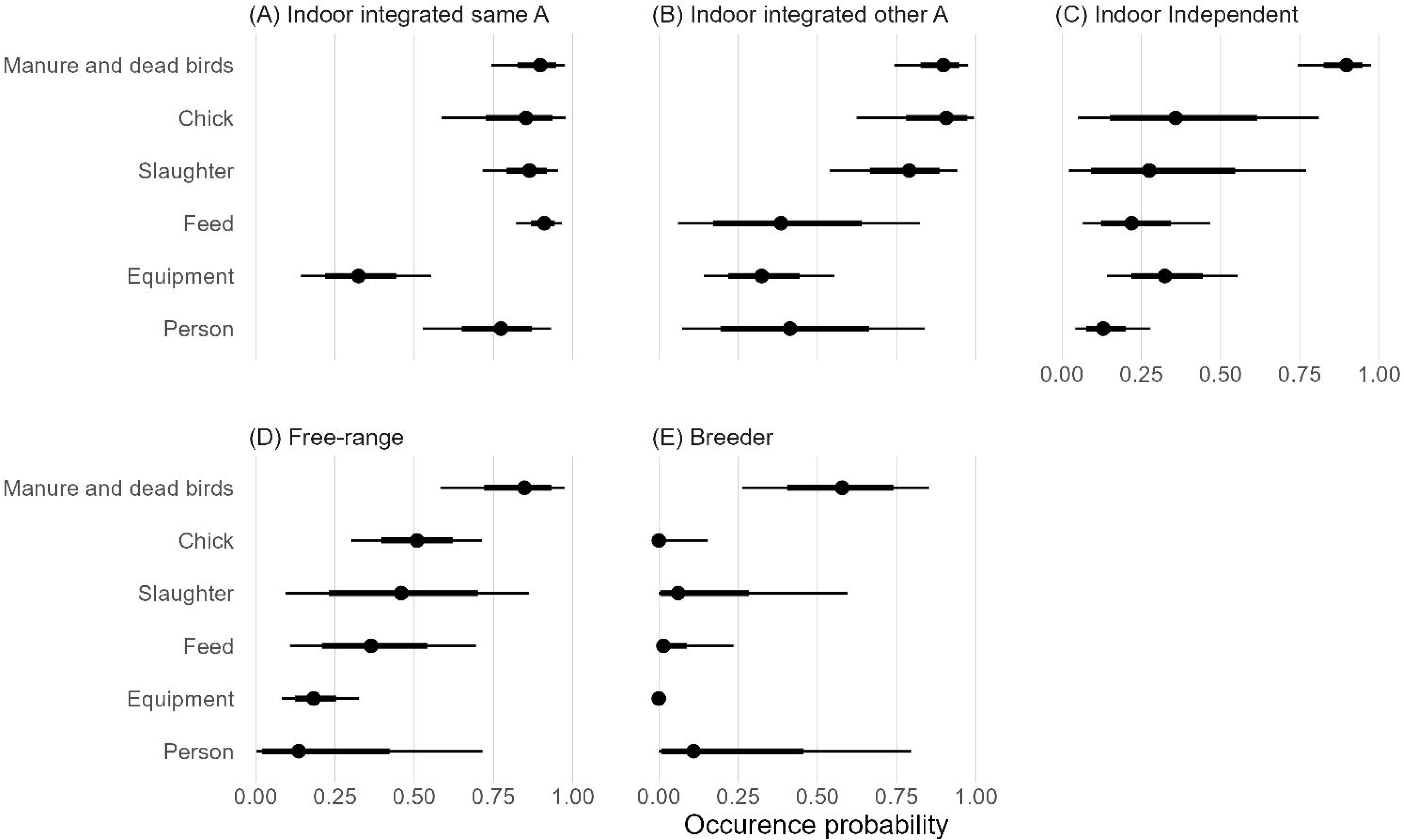
Probability (median, 50% and 80% quantile central intervals) of indirect contact in exposed integrated indoor broiler farms (free of infection) by pathway and type of source farm (infected): A) Indoor broiler farm integrated in the same farmer association, B) Indoor broiler farm integrated in another farmer association, C) Indoor broiler farm independent, D) Free-range broiler farm, E) Breeder farm.

### Farm biosecurity and pathways in the layer sector

Biosecurity levels were estimated to range from good to very good in indoor layer farms and from low to good in free-range farms, regardless of integration level. Pullet farms were considered with a good biosecurity level (Supplementary file 4). Due to their production, layer birds transport occurs at a lower frequency than broiler production. The pathways with the highest transmission probabilities were for both egg transport, person movement and shared equipment. Egg transport had the highest occurrence frequency in both indoor and free-range layer farms (median values of 45% and 30% respectively). Person movements and shared equipment were just as frequent as the other pathways but they are both associated with direct contact with birds.

### Farm biosecurity and pathways in selection farms

For both broiler and layer sectors, experts considered the biosecurity levels of selection farms to have high to very high biosecurity levels (Supplementary file 4), implying very low transmission probabilities, except for person movement. While breeder farm results were similar across sectors, egg transport and manure management appeared to have higher transmission probabilities in broiler than in layer sector (Supplementary file 3 Figure 3). If the transmission probabilities are very low in selection farms at high and very high biosecurity levels, they are closed to those observed in production farms for pathways such as person movement, transport of feed and eggs, and manure management if the biosecurity level of the farms reached a good level.

## Discussion

This study analysed the risk of virus transmission within French production networks, i.e. the risk that a virus released by an infected farm is introduced into a free farm across all possible indirect pathways and farm types. Most AI virus risk assessments have focused on risk factors related to the production network and to wild birds [28,29]. Some studies addressed the risk of introduction or release of AI viruses in farms with various production types [14,15,18,21], and a few explored the risk of virus transmission between farms types [18,30]. However, none considered the integration level of the farm or the variation of transmission risk across farm types.

This study used scenario trees model and a combination of literature data and experts’ elicitation partly based on the DELPHI method [31]. Unlike classic DELPHI method (reaching consensus) we used the difference of opinion among experts to estimate uncertainty about transmission probabilities. Hence, our study is subject to potential biases. To reduce the bias, we implemented six different tools: (1) different questionnaires to segment topics, (2) expert confidence level included into the statistical analysis, (3) option for experts to skip questions, (4) a comment section for each group of questions, (5) a guide to help the experts understand the questions and (6) a biosecurity grid to harmonize expert understanding. The expert elicitation was conducted in 2020. Even if the number of worldwide outbreaks increased drastically since 2021 [32] the experts were already aware of the threat because of 2015-16 and 2017-18 epizootics.

A key innovation of this study is the inclusion of biosecurity levels in the risk assessment. While biosecurity classifications developed for poultry farms generally provide a farm-level biosecurity score [26,33,34], few studies have assessed level of biosecurity by farm types (breeder, broiler or layer) [35] or integration level in the network [36,37] and, none have used this classification to assess the risk of disease transmission within a given farm type. Using an approach based on farm biosecurity enabled us to study a larger number of farm types and to generalize our results to any French poultry farm. The farm biosecurity level of an exposed farm (i.e. a farm that is free of infection) has a major impact on the risk of transmission of the AI virus between farms regardless of the pathway and of the biosecurity level of the source farm (i.e. a farm that is already infected), as previously highlighted in several studies [38–41]. In our study, farms dedicated to producing birds destined for reproduction (future breeders and breeder farms) mainly received a score of 5, indicating very high biosecurity, while production farms were scored 3, indicating good biosecurity. Fewer than 5% of the production farms were considered to have a very high level of biosecurity, in line with the results of a study conducted in New Zealand [33]. Another study in South Korea that measured the risk of HPAI for farms linked by livestock transport vehicles reported a low risk for breeder chicken farms [18]. In the present study the proportion of broiler farms with a low biosecurity level (level 2) was higher among independent farms than in integrated farms. Nonetheless, the differences observed need to be interpreted with caution as expert confidence was higher for integrated farms than for independent farms. Similar results were observed in layer sector except for independent indoor farms. However, only three experts out of five provided data for this type of farm (mean confidence of 3.7 out of 5) illustrating the lack of knowledge concerning this minority farm type. A French study provided information on biosecurity as a function of farm type (including broiler and layer) and location, but did not investigate independent farms [42].

This study highlighted that farm network organisation increases the occurrence probability and frequency of transmission pathways between farms within this network and thus the probability of viral transmission. First, this probability was highest between broiler farms that belong to the same farmers’ association. The probability of contact was highest between these farms, particularly via contact pathways such as people in direct contact with birds and feed transport trucks. This reflects the French poultry network in France where poultry farmers’ associations share feed manufacturer, technicians, and often veterinarians. The role of the poultry network in the spread of AI between farms was already highlighted in previous studies [11,13,16]. Second, egg transport plays a major role in the probability of transmission between layer farms [18,30]. Third, the existence of many of pathways between breeder farms increases the risk of transmission of the virus. Indeed, our results show that if the biosecurity level of a breeder farm that is an exposed farm is only ‘good’, the probability of transmission is similar to that of production farms. This result was confirmed by previous studies [35,38]. A study of HPAI H5 outbreaks in France between 2020 and 2022 showed that at least 9% of outbreaks occurred in a breeder farm [2].

The study focused only on indirect transmission pathways (i.e., humans or fomites) rather than direct pathways (i.e., live domestic birds). Live domestic bird movement is known to present high risk of virus transmission, especially in the duck sector [8]. However, this particular pathway is absent in broiler farms and only used at very infrequent intervals in layer and breeder farms (about once every 12-13 months). Improved truck cleaning and disinfection process would considerably reduce the risk of viral transmission within the network of farms. French studies detected the presence of the viral genome in half (in 2017) and in one third (in 2021) of trucks tested after disinfection [43,44]. In the present study, fomites management (i.e. a person, truck or equipment) outside farms was considered similar across farm types and insufficient to affect risk transmission. Disinfection of fomites inside the farm is included in the biosecurity classification. Based on poultry production experts, we considered that the frequency of occurrence of shared equipment was the same for all farms (once a month), regardless of the type or the network of farms, as this occurrence depends on management and location of the farm. While this impacts the relative role of shared equipment to viral transmission in farms with a low biosecurity level especially, it does not affect inter-farm comparisons.

Our study focused on the transmission pathways that have been identified as being the most at risk of transmission in the literature [10], but other transmission pathways, such as wind or vermin (i.e. mechanical vectors such as rodents or insects) are also mentioned. In France air-borne transmission is considered limited [45], even though some viral genome have been detected in air and dust in infected farms [46,47]. Other studies conducted in the USA suggest that airborne transmission might play a role in the spread of AI viruses [48,49]. Virus transmission through the movement of vermin between farms is possible but difficult to distinguish from other environmental factors [28,30,50]. These last pathways are often linked to farm proximity. High poultry density is known to have an impact in the spread of AI viruses [51]. Our study assumed that environmental transmission is constant across farm types regardless of their level of biosecurity. While this impacts geographic differentiation, future work could integrate spatial data.

Finally, this work provides a complete picture of the risk of AI viruses spreading within French broiler and layer production networks including the risk of transmission of the virus between any type of farms. To mitigate this risk, management measures should focus on biosecurity and controlling very frequent movements, especially of people, of feed, manure and dead birds. Network managers should be aware of intra-association movements and take the necessary actions, such as cleaning and disinfection, between visits to farms, to reduce this risk. The risk-based farm typology developed for this study can inform control measures such as restriction of movement or vaccination.

## Acknowledgements

The authors gratefully acknowledge all the experts who answered the questionnaire and the reviewers for their constructive comments. A preprint version of this article has been peer-reviewed and recommended by PCI Anim Sci (https://doi.org/10.24072/pci.animsci.100347).

## Data, scripts, code, and supplementary information availability

Data and scripts are available online: https://doi.org/10.18167/DVN1/MT1A1Z.

Supplementary material associated with this article is available online: https://doi.org/10.1101/2024.09.11.612235.

