## Supplementary file 2 for "Comparative risk-assessment of highly pathogenic avian influenza H5 viruses spread in French broiler and layer sectors"

Supplementary file 2 - Estimating the biosecurity level of French poultry farms

In order to classify French poultry farms according to their biosecurity level, we have defined a grid (Table 1). This grid lists all the criteria a farm must meet to be classified in a biosecurity level.

We use the definitions from the Order of February 8, 2016 on "biosecurity measures applicable in poultry and other captive bird farms as part of the prevention against avian influenza":

- "Farm": any agricultural facility in which poultry or other captive birds are reared or kept.
- “Production unit": any part of a farm which is completely independent of any other unit of the same establishment as regards its location and the routine management activities of the poultry or other captive birds kept there;

Note: For a poultry farm in confinement (building only, no outdoor run), a production unit corresponds to a building. For a free-range poultry farm (presence of an outdoor run) and for fattened palmipeds, a production unit corresponds to one or more pairs of buildings with runs (one or more runs for a building, but never a single run for several buildings).

- “Professional zone": area of the farm delimited outside the rearing zone, reserved for the circulation of authorized persons and vehicles, and for the storage or transit of incoming and outgoing products;
- “Farming area": area of the farm comprising all production units;
- “Farm site": area of the farm made up of the breeding zone and the professional zone;

For a farm to be classified in a biosafety level, it must comply with:

- All the criteria of the lower biosafety levels;

- All the criteria of the targeted level;

- Does not meet at least one of the criteria of the higher level.

Example: a blue ostrich farm that meets all the criteria of level 2, in which the sanitary lock is delimited and the farmer washes his hands on entry but does not change his boots, is classified as level 2.

Table 1: Criteria grid for different biosafety levels (from very low to very high)

| Biosecurity level | Criteria |
|---|---|
| 1<br>Very low | None |
| 2<br>Low | - The production unit comprises at least one building in which birds can be kept in confinement if necessary (including outdoor production farms in which birds can be kept in confinement, for example in the event of a health crisis).<br>- Presence of a sanitary airlock at the entrance to each production unit.<br>- Physical separation of the rearing area (fence, hedge, etc.).<br>- Protection of feed storage area from wild birds. |
| 3<br>Good | - Division of airlock into 2 or 3 zones using a board or equivalent.<br>- Change of boots or use of overshoes in the airlock<br>- Cleaning and disinfection of hands in the airlock<br>- Site effectively closed by a gate or chain (only persons authorized to work on the site may enter) |

| | |
|---|---|
| | - Check that vehicles and shared equipment are clean when entering the breeding site<br>- Implement a pest control plan (e.g. rodents) and restrict the entry of wild birds into the building (screened doors, windows and air inlets).<br>- For free-range farms, feeding and drinking points inside the building |
| 4<br>High | - Entire farm site physically separated (fence, hedge, etc.)<br>- Vehicles and shared equipment can be disinfected at the entrance to the farm.<br>- For free-range farms, runs protected by nets |
| 5<br>Very high | - Change of clothing and shower for personnel at the entrance to the site<br>- A decontamination area designated by a sign at the entrance to the site, with systematic disinfection of vehicles and equipment shared in this area.<br>- Sealing of building(s) against pests and wild birds |

Sources : [1–3]
