## Supplementary file 3 for "Comparative risk-assessment of highly pathogenic avian influenza H5 viruses spread in French broiler and layer sectors"

Supplementary file 3 – Results of the sensitivity analysis

Figures 1 and 2 display the results of the sensitivity analysis conducted on the type of farm with the lowest transmission probability in the broiler sector (i.e. future breeder farm).

The figure 3 illustrates the results of the sensitivity analysis on the biosecurity level of the source farm for breeder farms in broiler and layer sectors.

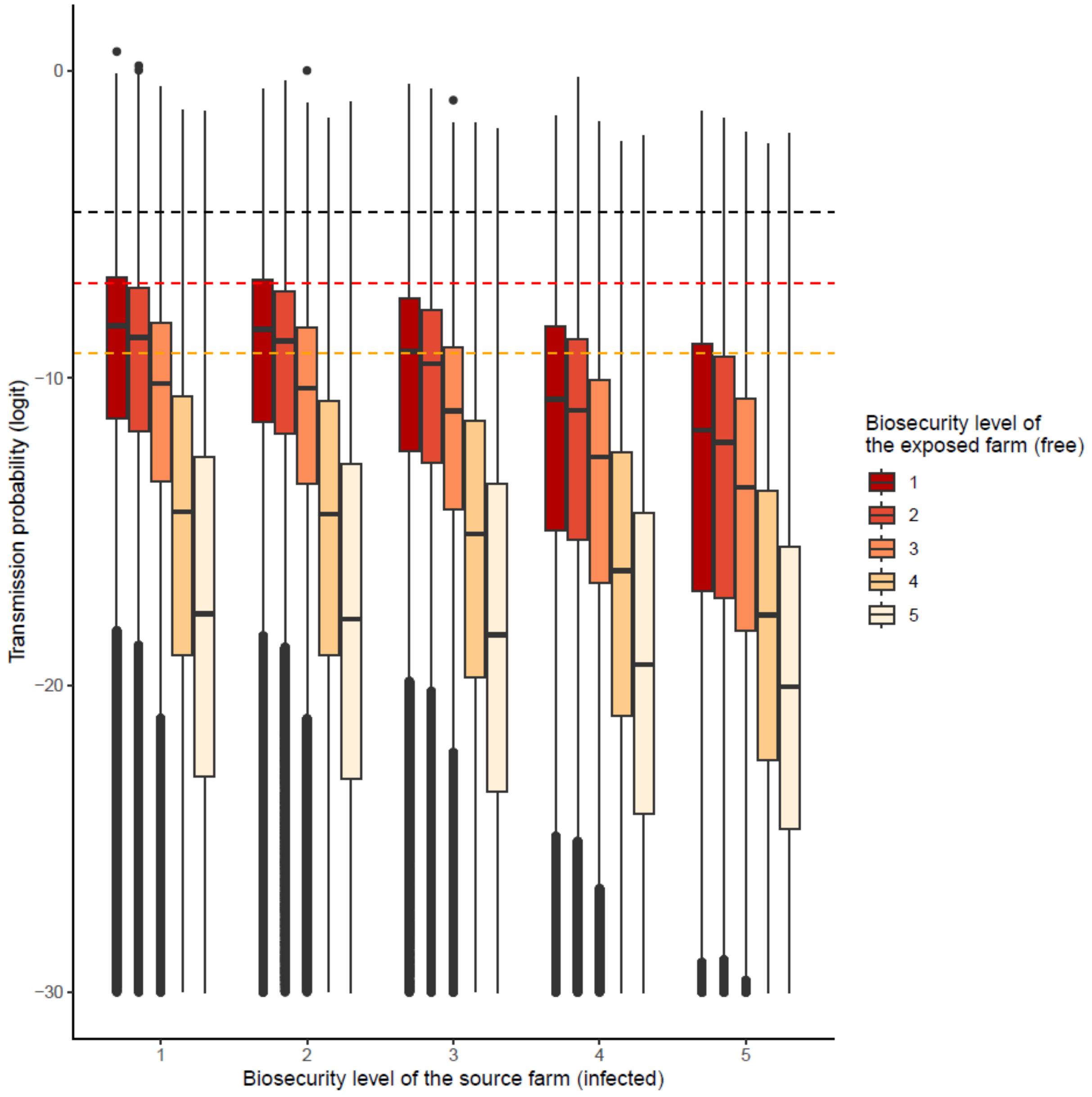


Figure 1: Distribution of the transmission probabilities across all **pathways** according to the biosecurity level of the source farm (x axis) and to the biosecurity level of the exposed farm (colours) in future breeder farm (broiler sector). Biosecurity level: 1 (very low), 2 (low), 3 (good), 4 (high) and 5 (very high). Threshold at 1/100 (black), 1/1000 (red) and 1/10 000 (orange).

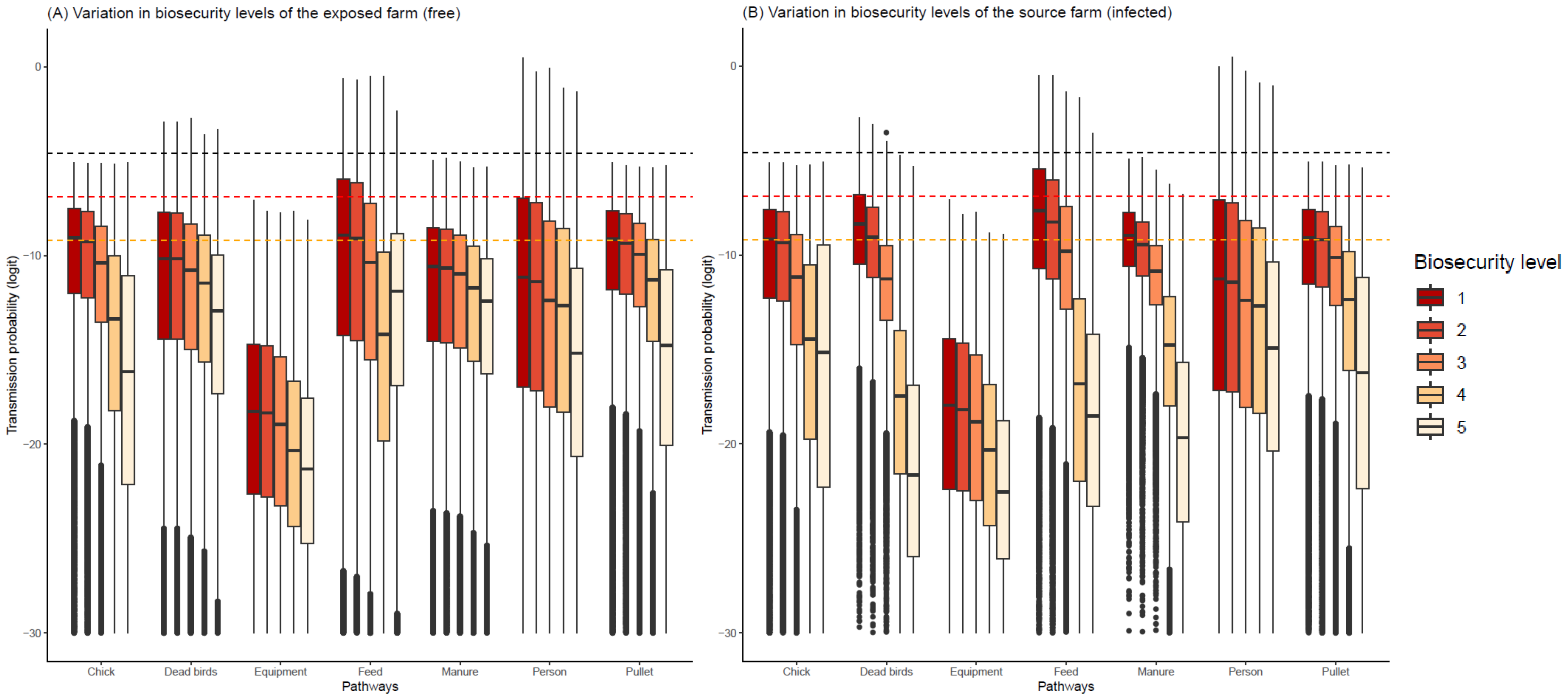


Figure 2: Distribution of the transmission probabilities in future breeder farm (broiler sector) as source and exposed farm (A) across all **biosecurity levels of the exposed farm (free)** according to the pathways (x axis) and to the biosecurity level of the source farm (infected) (colours), (B) across all **biosecurity level of the source farm (infected)** according to the pathways (x axis) and to the biosecurity level of the exposed farm (free) (colours). Biosecurity level: 1 (very low), 2 (low), 3 (good), 4 (high) and 5 (very high). Threshold at 1/100 (black), 1/1000 (red) and 1/10 000 (orange).

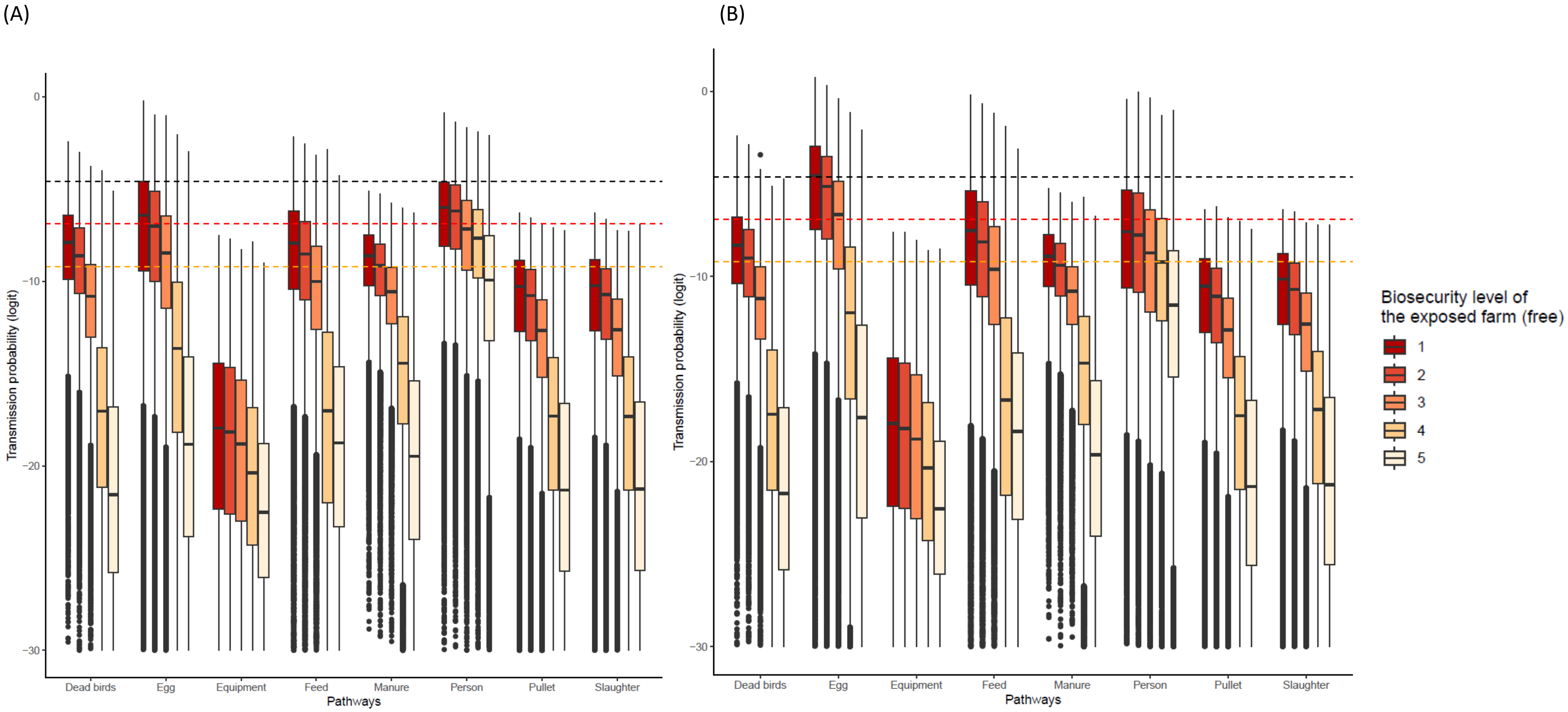


Figure 3: Distribution of transmission probabilities across all **biosecurity levels of the source farm (infected)** according to the pathways (x axis) and to the biosecurity level of the exposed farm (free) (colours). Biosecurity level: 1 (very low), 2 (low), 3 (good), 4 (high) and 5 (very high). Threshold at 1/100 (black), 1/1000 (red) and 1/10 000 (orange). A: breeder farms for layer production, B: breeder farms for broiler production
