## Supplementary file 4 for "Comparative risk-assessment of highly pathogenic avian influenza H5 viruses spread in French broiler and layer sectors"

Supplementary file 4 – Biosecurity levels

### Broiler sector

Figure 1 displays the estimated distribution of biosecurity levels provided by the experts on the broiler sector (7 experts) for the six different types of farms (integrated and independent in-door broiler farms, integrated and independent free range broiler farms, pullet future breeder farms, and breeder farms). The majority of integrated broiler farms were scored with a good biosecurity level, with 53% of in-door and 45% of free range estimated at level 3 (Figure 1). The distribution of biosecurity levels was very similar for the two types of integrated farms (indoor and free-range), while independent indoor broiler farms were scored with a higher biosecurity level than independent free-range broiler farms, i.e. the majority of indoor farms at a good level (43%) and of free-range farms at low level (42%). For both indoor and free-range farms, independent broiler farms were estimated to have a lower biosecurity level than integrated farms, as the majority of farms were considered to have a low biosecurity level (level 2). Future breeder and breeder farms were scored with the highest biosecurity level as a majority of farms were considered to have a very high biosecurity level (respectively 67% and 66%) (level 5). The experts were generally very confident in their answers on integrated farms (indoor and free-range) and selection farms while they were moderately confident in their answers on independent farms (indoor and free-range).

A- Integrated indoor broiler farms

B- Independent indoor broiler farms

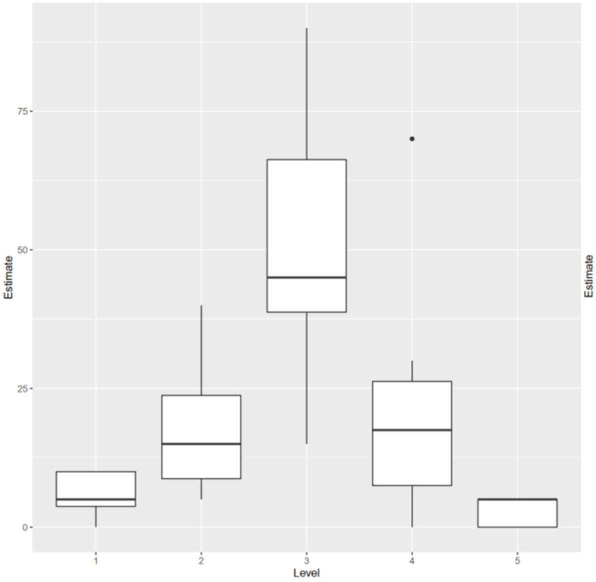


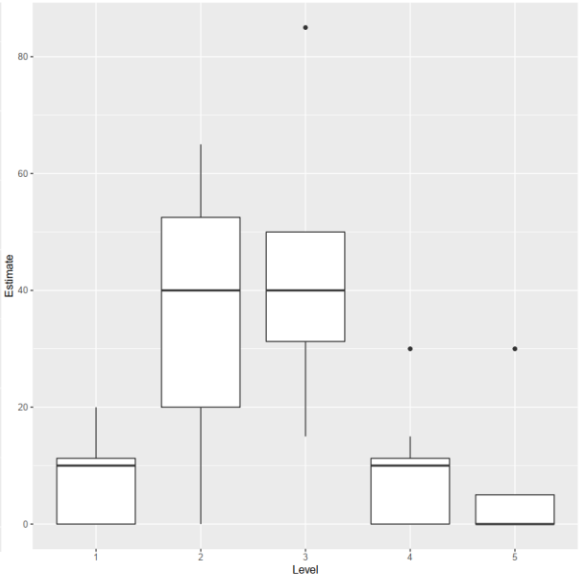


C- Integrated free-range broiler farms

D- Independent free-range broiler farms

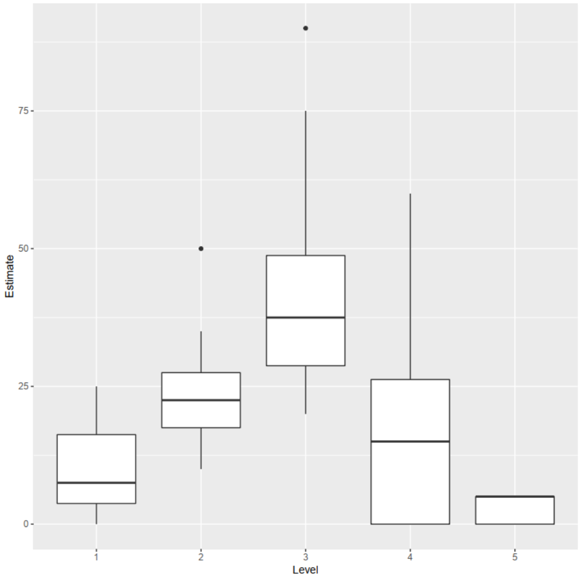


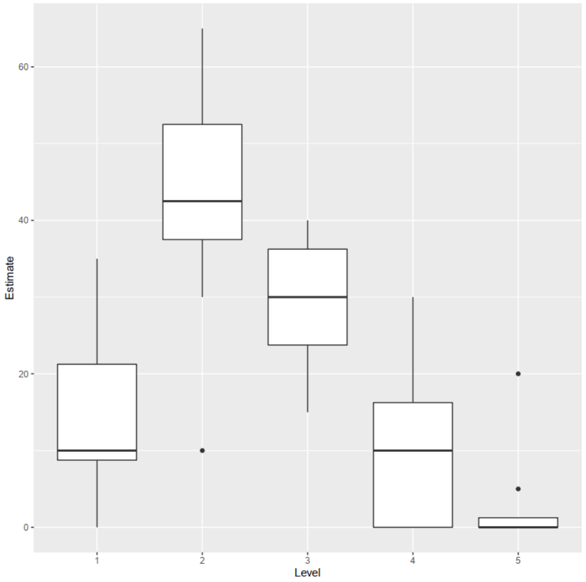


E- Future breeder farms

F- Breeder farms

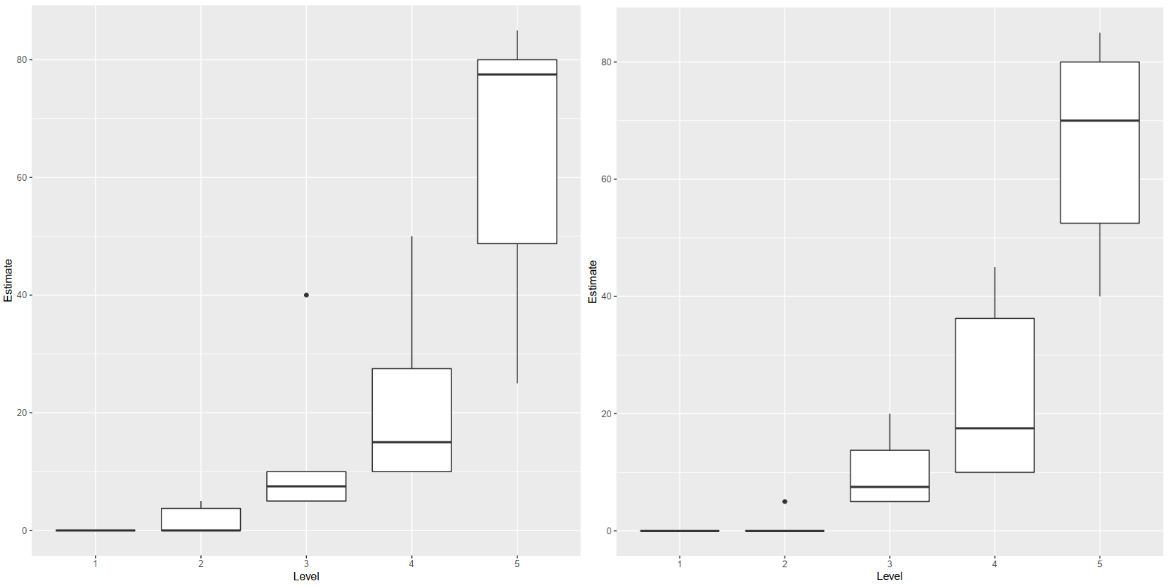


Figure 1: Distribution of farm biosecurity levels according to the 6 defined farm types (A- Integrated indoor broiler farms, B- Independent indoor broiler farms, C- Integrated free-range broiler farms, D- Independent free-range broiler farms, E- Future breeder farms, F- Breeder farms) (according to 7 experts in 2020). Figures axis: Level: Biosecurity levels: 1 (very low), 2 (low), 3 (good), 4(high) and 5 (very high); Estimate: Proportion of farms the experts attributed to each biosecurity level. Boxplot provides the minimum (end of the lower line), first quartile (far lower of the box), median (line in the centre of the box), third quartile (far higher of the box) and the maximum (end of the higher line). Extra dots are outliers.

Layer sector

Figure 2 displays the estimated distribution of biosecurity levels provided by the five experts on the layer sector for the seven farm types (future layer farms, integrated and independent in-door layer farms, integrated and independent free range layer farms, pullet future breeder farms, and breeder farms). Indoor layer farms were estimated to have between very good biosecurity level (40% of integrated and integrated farms) and good biosecurity level (38% integrated and 25% independent farms) (Figure 2). Free-range farms were estimated to have between good biosecurity level (35% integrated and 28% independent) and low biosecurity level (30% integrated and 35% independent). Pullet farms were considered with a good average biosecurity level (50%). As for the broiler production, breeder and future breeder farms were scored with a very good biosecurity level (75% and 78% respectively).

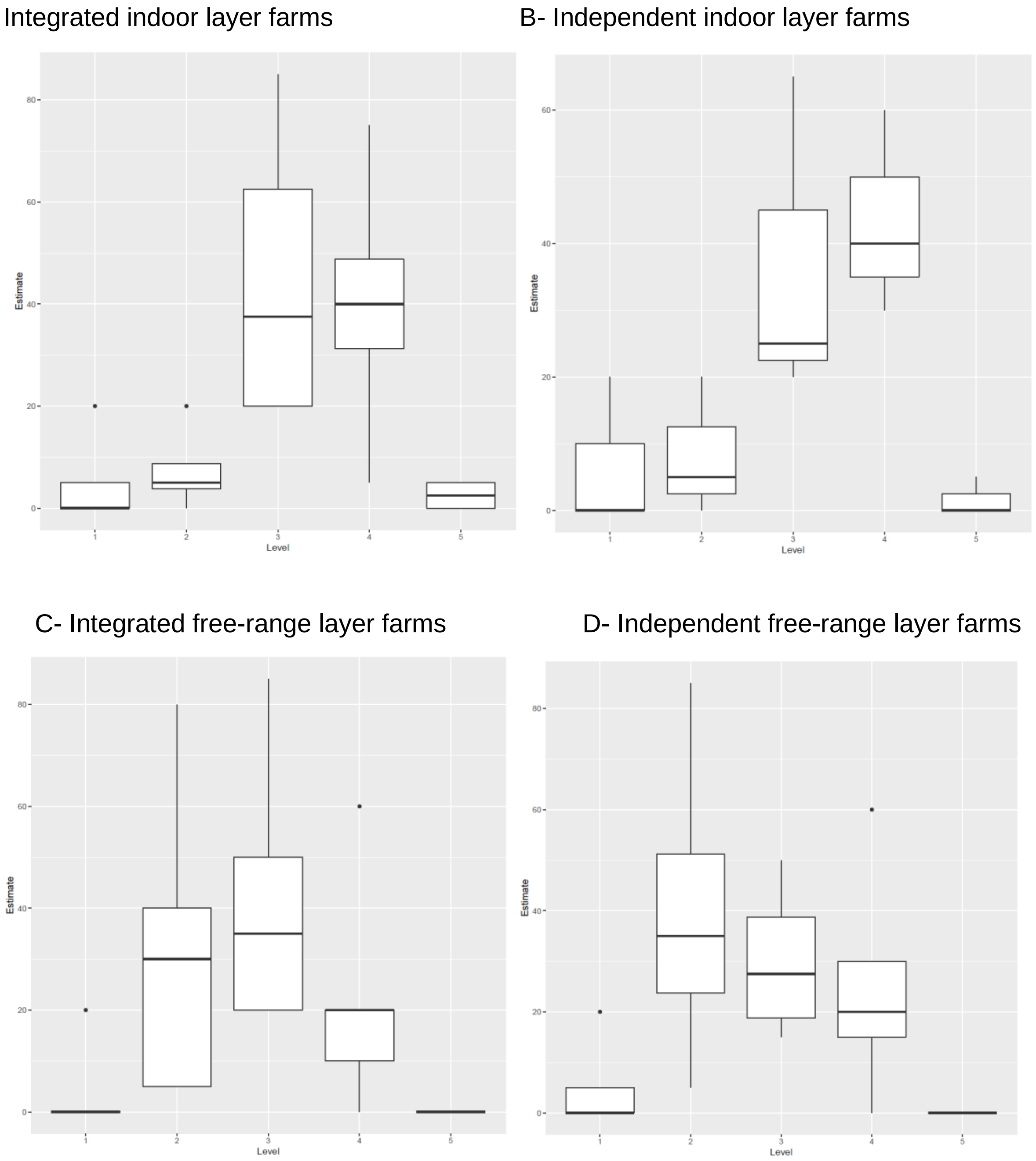

E- Future breeder farms

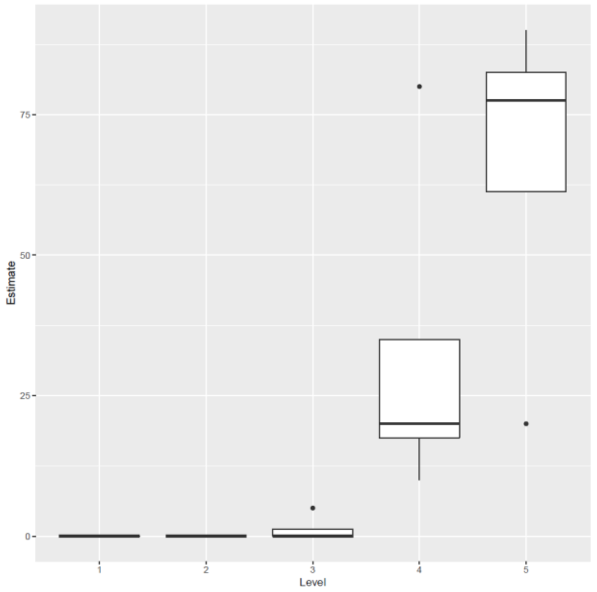


F- Breeder farms

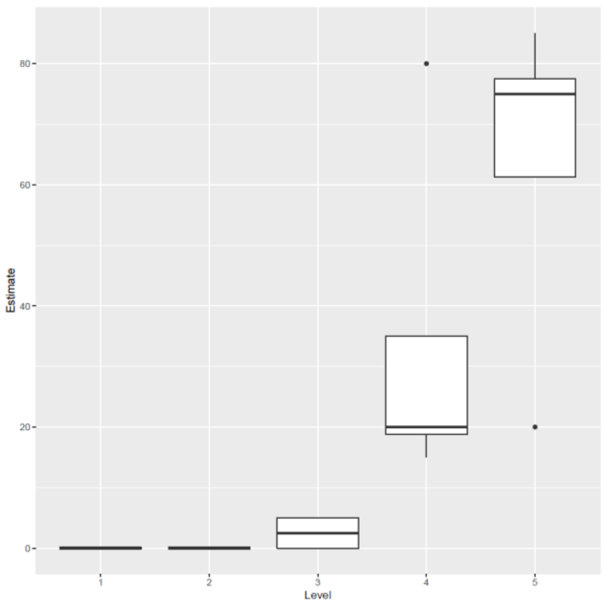


G- Pullet farms

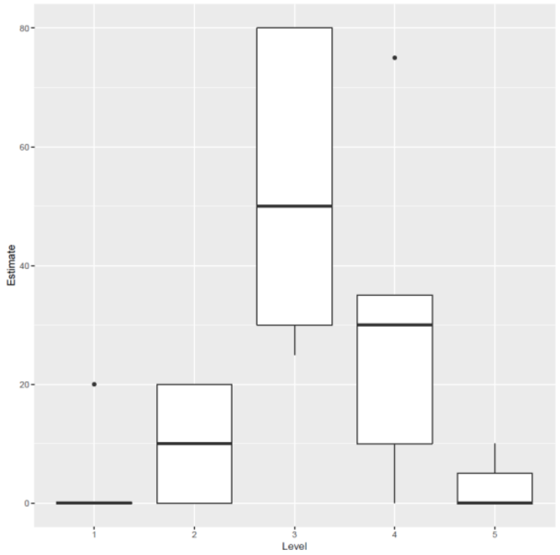


Figure 2: Distribution of the farm biosecurity level according to the different farm types in 2020 (A- Integrated indoor broiler farms (4 experts), B- Independent indoor broiler farms (3 experts), C- Integrated free-range broiler farms (5 experts), D- Independent free-range broiler farms (4 experts), E- Future breeder farms (4 experts), F- Breeder farms (4 experts), G- Pullet farms (5 experts)). Figures axis: Level: Biosecurity levels: 1 (very low), 2 (low), 3 (good), 4(high) and 5 (very high); Estimate: Proportion of farms

the experts attributed to each biosecurity level. Boxplot provides the minimum (end of the lower line), first quartile (far lower of the box), median (line in the centre of the box), third quartile (far higher of the box) and the maximum (end of the higher line). Extra dots are outliers.
